# Long-term control of *Salmonella* after transient T3SS-2 inhibition

**DOI:** 10.1101/2025.10.30.685576

**Authors:** Alaa Alhayek, Beatrice Claudi, Sila Nizam, Helen Atassi, Dirk Bumann

## Abstract

Anti-virulence approaches are promising alternatives for traditional antibiotics to control bacterial infections. Several inhibitors show impressive activities in animal infection models, but the relative contribution of specific virulence inhibition vs off-target effects on both the bacteria and host remain unclear. Here, we developed *Salmonella* with switchable virulence by putting the type 3 secretion system-2 (T3SS-2) which is essential for systemic virulence, under the control of doxycycline. In infected mice given low-dose doxycycline in drinking water, the strain showed normal fitness and virulence. Doxycycline withdrawal shut down T3SS-2, arrested *Salmonella* replication and resolved disease symptoms. After ten days of T3SS-2 inhibition, reintroducing doxycycline restored replication, but bacterial loads remained stable, indicating strengthened host immunity. These effects were comparable to treatment with fluoroquinolone antibiotics, a highly effective therapy for human systemic salmonellosis. Thus, selective T3SS-2 inhibition may offer a suitable alternative for controlling invasive *Salmonella* infections.

## Main

Bacterial infections remain a major threat to human health. Most pathogens have evolved resistance to available antibiotics ^1^. There has also been a long gap in antibiotic discovery, with few novel antibiotics coming ^2^. However, the increasing understanding of bacterial pathogenesis has revealed potential non-antibiotic approaches. One of these is virulence inhibition which disarms pathogens in the interaction with the host immune system rather than killing them. This minimizes collateral damage to the host microbiome and slows resistance development ^3,4^. This approach has a long history of success, from early toxin-neutralizing (*e*.*g*., tetanus) to modern monoclonal antibodies ^5,6^ and small-molecule inhibitors ^4^. Many of these small molecules show potent in vitro activity, but variable in vivo activity ^7–9^. Further development of the most promising leads also remains scarce. Open questions include: can virulence inhibitors clear infections on their own? Can such inhibitor lead to long-term control of infections without relapsing disease after termination of virulence inhibition?

One widely studied virulence system that is suitable to address these questions, is type III secretion system (T3SS), in which an injectosome translocates virulence factors from the bacterial cytosol into host cells ^10^. T3SS is essential for virulence of diverse Gram-negative pathogens, including *Pseudomonas aeruginosa, Shigella, Yersinia*, and *Salmonella* ^11,12^. Among these, *Salmonella enterica* is a major cause of mortality worldwide and becomes increasingly untreatable ^13,14^ motivating the WHO to classify it as a priority pathogen for antimicrobial discovery ^15^. *Salmonella* encodes two T3SSs: T3SS-1, which mediates epithelial invasion, and T3SS-2, which supports intracellular growth in macrophages ^16^. *Salmonella* mutants lacking T3SS-2 activity are avirulent for systemic disease in mice and humans ^17,18^. T3SS-2 has therefore emerged as a promising target for anti-virulence development. However, a large part of the effector proteins translocated through T3SS-2 have anti-inflammatory activities ^19^, so abrupt inhibition could exacerbate inflammation. T3SS-2 inhibitors have been reported ^7,20,21^. Some of them reduce mortality in mouse infection models ^22–25^ (and human urinary-tract infections ^26^), implying that T3SS-2 activity is required continuously throughout infection. However, some of the inhibitors are likely confounded with off-target effects both in the bacteria and in mammalian host cells ^22,25^. Thus, the immediate and long-term impact of transient T3SS-2 inhibition in highly infected mice remains unclear.

### Establishing and validating a switchable anti-virulence approach in *Salmonella*

To determine the impact of anti-virulence treatments on *Salmonella* control in vivo, we developed a switchable system that activates or inactivates T3SS-2 without affecting other aspects of *Salmonella* biology or the host (Fig. 1a). T3SS-2 depends on the transcription factor SsrB ^27^. We inactivated *ssrB* by multiple STOP codons across the gene to reduce polar effects on downstream *ssrA*. This abolished the activity of a SsrB-dependent *P*_sifB_-*gfp* transcriptional fusion encoding an unstable variant of GFP (GFP-OVA ^28,29^) in T3SS-2 inducing conditions ^30^. We complemented the *ssrB* mutant with a doxycycline (DOX)-inducible *P*_tetA_-*ssrB* expression cassette. We used DOX as inducer because of its bioavailability in mice ^31^ and stability in acidified drinking water ^32^. In presence of 0.125 mg/L DOX (well below the minimum inhibitory concentration, 1-2 mg/L), *P*_sifB_-*gfp* regained full activity (Fig. S1b). However, there was also substantial activity even in absence of DOX, suggesting leaky *ssrB* expression (Fig. S1b). We optimized the dynamic range of switchable *ssrB* using an inefficient ribosomal binding site, resulting in low baseline and full DOX-induced activity of *P*_sifB_-*gfp* (Fig. 1b). We also included a constitutive *mCherry* expression cassette (Fig. S1a) to detect all *Salmonella* regardless of *P*_tetA_-*ssrB* activity. We called this construct pVIR-ON/OFF.

**Figure 1.**
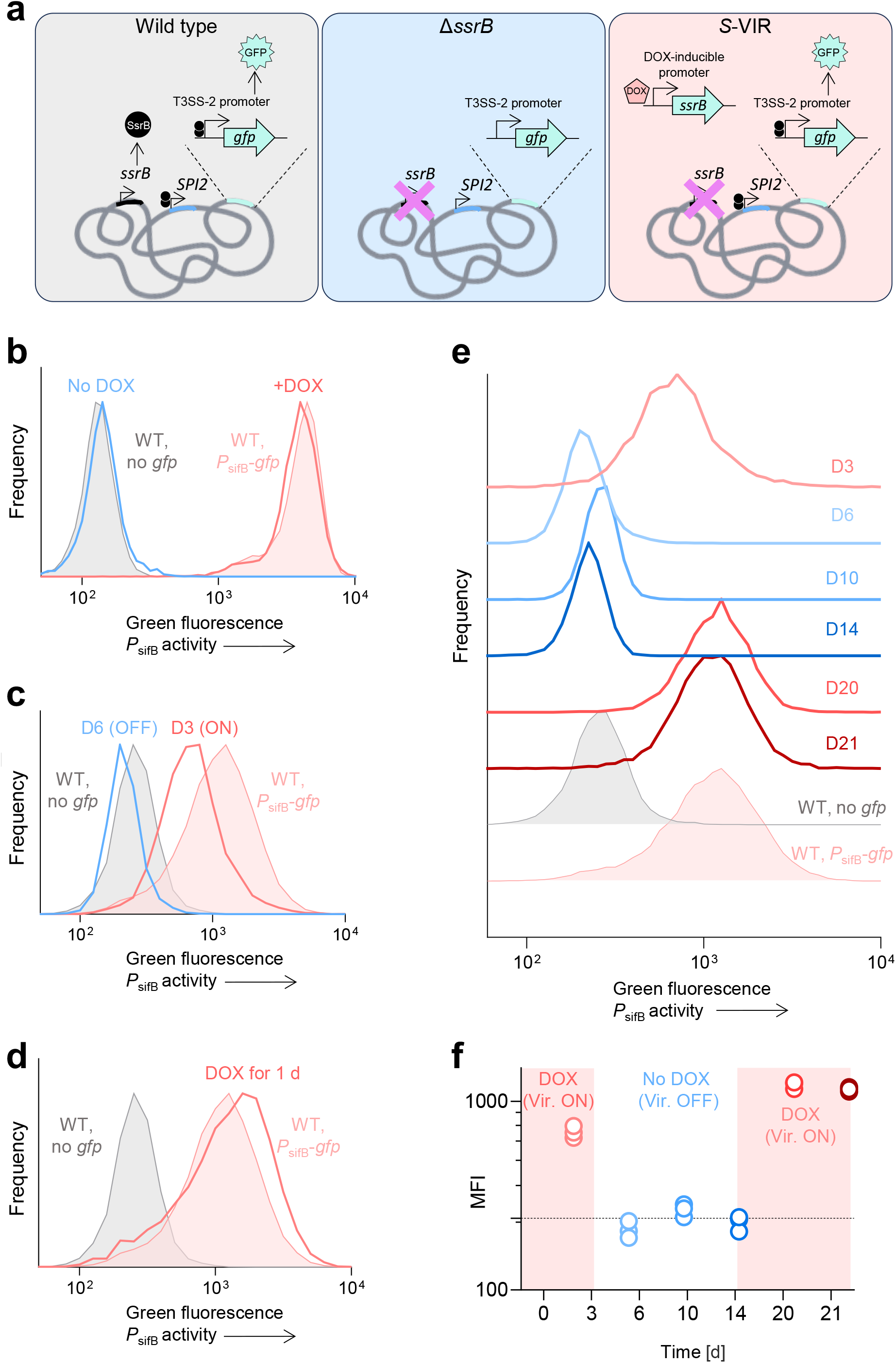
Validation of a switchable anti-virulence system in *Salmonella*. (a) Schematic description of the switchable system controlling T3SS-2. (b) In vitro validation of switchable anti-virulence, (DOX, Doxycycline). Autofluorescence of a (WT, wild-type) *Salmoenlla* (gray) and induced *P*_sifB_-*gfp* activity in WT is shown in (orange). Pooled data from two-independent experiments. (c) Green fluorescence of the switchable virulence *Salmoenlla* in mice receiving 0.01 mg/mL DOX in drinking water. Induced *PsifB-gfp* (orange) is high with DOX; after 3 days of DOX withdrawal, *PsifB-gfp* returns to baseline (blue). Autofluorescence of a (WT, wild-type) *Salmoenlla* (gray) and induced *P*_sifB_-*gfp* activity in WT is shown in (orange). Pooled data from three mice per group. (d) Green fluorescence after 6 days without DOX and one day of re-exposure. Autofluorescence of a (WT, wild-type) *Salmoenlla* (gray) and induced *P*_sifB_-*gfp* activity in WT is shown in (orange). (e) Time course of splenic GFP signal under DOX until day 3 and from day 14; pooled data from three independently infected mice. Autofluorescence of a (WT, wild-type) *Salmoenlla* (gray) and induced *P*_sifB_-*gfp* activity in WT is shown in (orange). Data for individual mice are shown in panel f. (f) Median green fluorescence intensity (MFI) for data shown in panel e. Each circle represents one mouse; the bar is the mean. The dashed line represents the MFI of wild-type *Salmonella* without *gfp*.

To test pVIR-ON/OFF in vivo, we provided DOX in drinking water to mice. We used mice with functional SLC11A1 to avoid sever immunosuppression ^33,34^. Previous studies used 0.15 to 2 mg/mL for inducing microbial gene expression from *P*_tetA_ ^32,35^. To minimize potential antimicrobial effects of DOX on *Salmonella*, we initially used 0.1 mg/mL DOX from 1d prior to and throughout infection. Mice infected with *Salmonella*/pVIR-ON/OFF (*S*-VIR) under these conditions developed symptoms comparable to mice infected with wild-type *Salmonella* without DOX. Activity of *P*_sifB_-*gfp* in spleen at d5 post infection were also comparable (Fig S1d). Conversely, infection of mice with *S*-VIR without DOX caused no disease symptoms (data not shown), little bacterial growth as reported before^36^, and only baseline *P*_sifB_-*gfp* activity similar to data for a *Salmonella ssrB* mutant (Fig S1c). Thus, *S*-VIR provided conditional SPI-2 activity, fitness, and virulence in vivo.

To simulate an anti-virulence treatment, we infected DOX-treated mice with *S*-VIR and withdrew DOX at 5d post-infection. However, at d6 we observed still high *P*_sifB_-*gfp* activity (Fig S1d), suggesting ongoing T3SS-2-activity. This could be due to slow elimination of DOX from tissue ^31^ and/or stability of previously made SsrB. We thus further reduced the DOX concentration to 0.01 mg/mL and withdrew DOX already at d3 post-infection. Under these conditions, disease symptoms and spleen *S*-VIR loads still increased until d6 indicating high fitness and virulence, but *P*_sifB_-*gfp* activity declined to baseline (Fig. 1e). When these mice were exposed again to 0.01 mg/mL DOX in the drinking water, *S*-VIR regained full activity (Fig. 1d & f). These data demonstrated that *S*-VIR was a fully functional switchable virulence construct.

### Disease relapse but CFU control post-anti-virulence

To determine the impact of virulence suppression and regain, we infected DOX-treated mice with *S*-VIR, withdrew DOX at d3 post-infection, and re-administered DOX from d14 modulating T3SS-2 activity accordingly (Fig 1f & 2a). Disease symptoms and bacterial loads in spleen increased until d6 (Fig 2a & b). In the absence of DOX, symptoms resolved and bacterial loads remained constant, confirming the suppression of virulence without eradicating the bacteria. Constant bacterial loads could reflect a replication arrest and no host-mediated killing, or a dynamic equilibrium between ongoing replication and killing. To distinguish between these alternatives, we combined *P*_tetA_-*ssrB* with *timer*^bac^ expression (Fig S1f) which reports bacterial replication rates *via* fluorescence spectral shifts ^34,37,38^. TIMER^bac^ revealed normal *Salmonella* replication at d3 in presence of DOX, but replication arrest at d10 and d14 in absence of DOX (Fig 2c). Thus, suppression of SPI-2 stopped replication in vivo without enabling host-mediated killing, consistent with the phenotype of SPI-2-deficient mutants ^36,39^.

**Figure 2.**
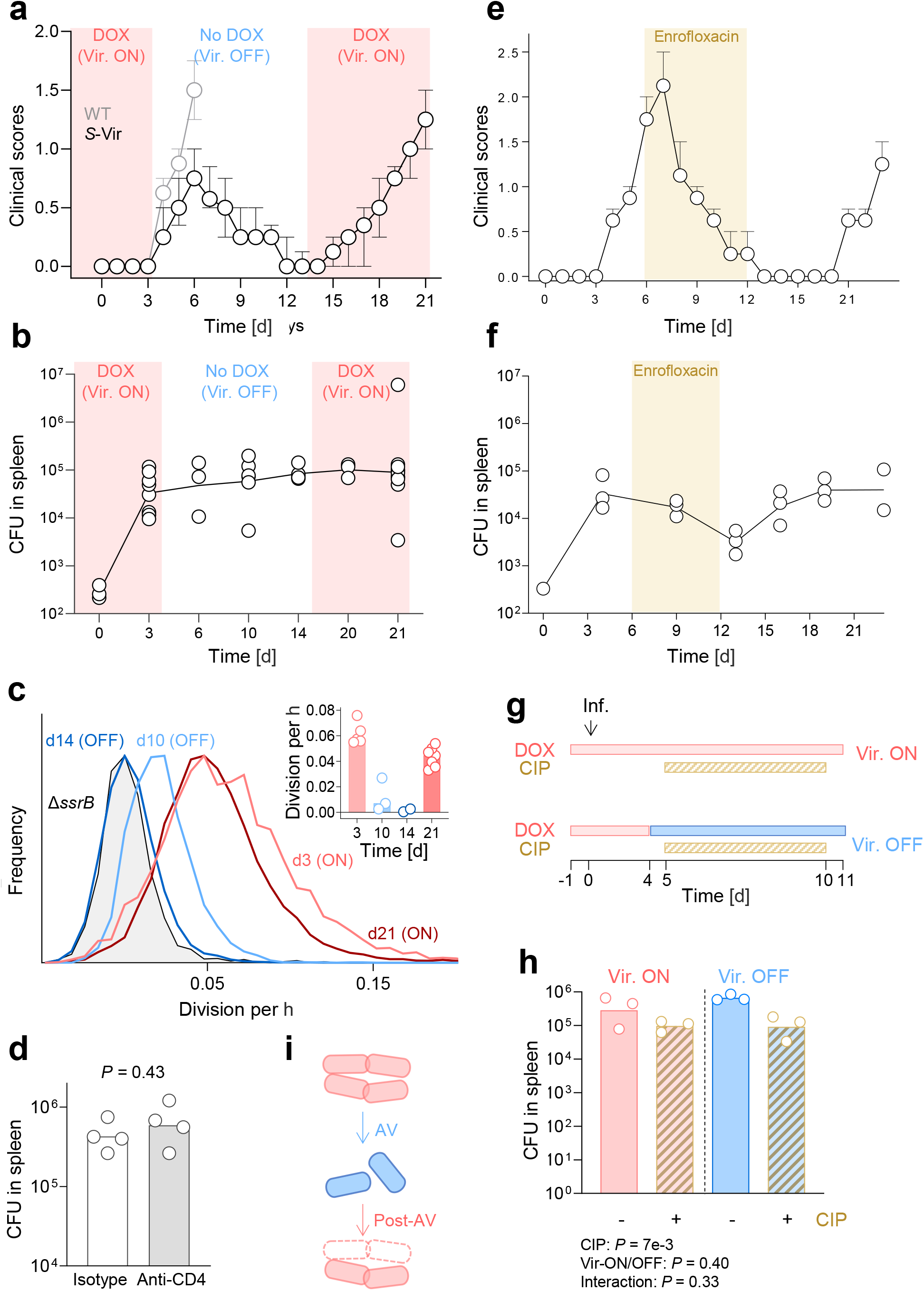
In vivo impact of virulence modulator. (a) Clinical scores and (b) spleen bacterial loads in mice infected with the switchable-virulence strain (S-VIR). (DOX, Doxycycline) was given from day 1 to day 3, withdrawn until day 14, then re-administered until day 21, (WT, wild-type). *(c) Salmonella* division rates. The histograms represent pooled three to seven mice. Data for each individual mice are shown in the inset. Each circle represents one mouse; the bars show the mean. *(d) Salmonella* burdens in spleen in mice anti-CD4–treated versus isotype control mice. Each circle represents an individual mouse; the bars show the geometric means. (e–f) Effects of enrofloxacin treatment on clinical scores (e) and spleen CFU (f). (g) Timeline of combined treatment with ciprofloxacin (CIP) and DOX to modulate T3SS-2 activity. (h) Comparison of antibiotic alone versus combined treatment (CIP + T3SS-2 OFF). Each circle represents an individual mouse; the line shows the geometric mean. (i) Proposed working model for anti-virulence treatment. Post anti-virulence replication restarted, but host killing leads to partial control.

Upon re-administration of DOX from d14, restarted of *Salmonella* replication (Fig. 2b & 2c). However, bacterial loads remained constant, suggesting enhanced *Salmonella* killing by the host. These enhanced host activities were associated with re-appearance of increasing symptoms. Thus, virulence suppression mitigated disease symptoms and stopped *Salmonella* growth. During this time, the host gained the capability to control *Salmonella* net growth even after restoring virulence, but this was insufficient to eradicate the bacteria and to prevent relapsing disease (Fig 2i).

### Minimal CD4^+^ T-cell role in post–T3SS-2 protection

CD4^+^ T-cells contributes to protective immunity against *Salmonella* after vaccination or during chronic infection ^40–43^. To test their contribution to post–anti-T3SS-2 control, mice received depleting anti-mouse CD4 or isotype control antibodies on d 14, 16, and 18 after termination of virulence inhibition. We used 800 µg per mouse, a dose reported to achieve efficient depletion ^44^. *Salmonella* burdens were comparable in anti-CD4–treated mice *vs*. controls (Fig 2d), indicating limited impact of CD4^+^ T-cell. Thus, alternative mechanisms such as trained innate immunity may contribute^45^. This requires further investigations.

### Disease relapse but CFU control after enrofloxacin antibiotic

To Compare anti-T3SS2 effect with clinically potent anti-*Salmonella* antibiotics. We treated infected mice with enrofloxacin, the veterinary prodrug of ciprofloxacin. Enrofloxacin cleared only ∼1 log CFU but also permitted control of bacterial loads after treatment albeit with relapsing disease symptoms (Fig 2 e & f).

### No synergy between T3SS-2 switch-off and antibiotic treatment

It has been proposed that T3SS-2 activity is essential for intracellular *Salmonella* survival during antibiotic exposure ^46^. Switching off T3SS-2 could thus sensitize *Salmonella* to antimicrobial chemotherapy. To test this, we treated infected mice from day 5 with the widely used fluoroquinolone ciprofloxacin using a dosing regimen as in human patients (10 mg/kg body weight, q12 h for 5 days) (Fig 2g). This treatment showed moderate efficacy (∼1 log reduction in splenic CFU), consistent with the limited reported efficacy against *Salmonella* in mice carrying functional SLC11A1 ^30^ (Fig. 2e–h). Maintaining T3SS-2 in the OFF state throughout therapy did not further reduce bacterial loads or clinical signs. These findings align with prior work showing only additive, rather than synergistic, effects of an SsrB-targeting anti-virulence compound with colistin ^22^. The results were also consistent with unaltered antibiotic survival of T3SS-2–deficient *Salmonella* ^18,37^. Together, the data indicate that T3SS-2 activity is dispensable for *Salmonella* antibiotic tolerance even at advanced disease stages and that combining T3SS-2 inhibition with standard antibiotics is unlikely to yield synergy.

## Discussion

Targeting virulence rather than directly killing bacteria is a promising strategy to control infection, offering more druggable targets, reduced selection for resistance, and less disruption of the microbiota ^3,4^. However, anti-virulence compounds are not supposed to kill the bacteria directly ^3,4^, posing a risk of bacterial persistence and relapsing disease after treatment termination. Such disarmed bacteria might be easier to clear by host immunity, enabling long-term infection control. Some anti-virulence compounds show impressive bacterial clearance in animal infection models ^22–24^ and partial effects in human patients ^26^. Interpretation of such results remain challenging due to documented off-target effects in both bacteria and host cells.

To assess the impact of specific targeting a single virulence system without perturbing bacterial or host physiology, we engineered a genetic system that enabled us to switch on and off virulence of *Salmonella* in infected mice using simple changes in the drinking water. We targeted T3SS-2 which is essential for *Salmonella* virulence ^17,18^ and exploited as target for two advanced anti-virulence compounds ^22,26^. We inactivated the entire T3SS-2 by mutating the central transcription factor *ssrB*, and then complemented the strain with a DOX-inducible active *ssrB* allele. We showed that low DOX concentrations and without detectable effects in mice, fully restored virulence. Switching off T3SS-2 in infected mice arrested bacterial replication and – with some delay^47^ – alleviated disease symptoms. Importantly, this transient, on-demand inhibition is more relevant to anti-virulence therapy compared to well-characterized permanently T3SS-2–deficient *Salmonella* mutants, which replicate poorly and cause no detectable disease symptoms and inflammation ^18,37^.

Switching off T3SS-2 for ten days did not clear infection, consistent with the long-term in vivo survival of T3SS-2 mutants and indicating that host immunity alone cannot eliminate disarmed *Salmonella*. These data also imply that prior reports of eradication by T3SS-2 inhibitors likely reflect off-target antibacterial or host effects ^22,25^. Conversely, we did not observe escalating inflammation, suggesting that the anti-inflammatory functions of T3SS-2 – which might be important for shaping *Salmonella* microenvironments – exert limited effects at the organismal level. This supports the safety of T3SS-2 inhibition.

When T3SS-2 was reactivated, *Salmonella* resumed replication, yet total bacterial loads remained stable, indicating that host immunity balanced *Salmonella* proliferation with killing. This differed markedly from the initial infection phase when active T3SS-2 drove exponential bacterial proliferation. Thus, transient T3SS-2 inhibition allowed the host to develop partially protective immunity. CD4^+^ T-cell depletion did not diminish this partial protection, suggesting alternative mechanisms that will require further investigation. Nevertheless, this partial control was insufficient to prevent relapse of disease symptoms.

Comparison with clinically potent anti-*Salmonella* antibiotics revealed slight better, but again incomplete clearance and similar control of bacterial loads followed by relapse after treatment cessation. Combining T3SS-2 inhibition with antibiotics did not improve clearance. Together with prior in vivo data, these findings do not support a critical role for T3SS-2 in *Salmonella* survival during antibiotic exposure, contrary to proposals based on cell-culture studies ^46^.

T3SS-2 inhibition seemed as effective as standard antibiotics at suppressing *Salmonella* regrowth, suggesting that this approach could represent a suitable alternative. However, neither strategy prevented relapse in mice. Importantly, similar antibiotic regimens in human systemic salmonellosis achieve high cure rates and prevent relapse, indicating a limited predictive value of commonly mouse model. This could be related to stronger innate immunity / less virulent *Salmonella* strains in human patients compared to mouse infections. We used mice with restored SLC11A1 function which increases their anti-*Salmonella* resistance, but all infected animals would ultimately succumb, whereas human mortality is 10–30%, confirming lower clinical virulence. Taken together, these observations support prior concerns that standard mouse models are overly stringent for preclinical evaluation ^48^ and suggest that strains with better control of invasive *Salmonella* may more accurately predict the clinical utility of anti-virulence approaches ^49–51^.

In conclusion, we established a versatile model for transient on-target virulence inhibition in *Salmonella*. Selective inhibition of T3SS-2 did not eradicate infection, but it enabled post-treatment control, with efficacy approaching that of antibiotics that are highly effective against human systemic salmonellosis.

## Materials and Methods

### Bacterial strains and plasmids

*Salmonella* strains used in this study were based on SME51, a prototrophic *hisG*^Leu69^ derivative of *Salmonella enterica* serovar Typhimurium SL1344 ^39,52,53^. Reporter strains carried inactivated *ssrB* by nonsense (STOP) mutations across the gene ^54^ (chromosomal coordinates in SL1344 GenBank FQ312003.1: 1,433,019-1433657, M207X, M189X, G164X, E139X, M91X, M78X, G61X, M20X,

M1X). Also, the strain carried low-copy episomal pSC101 derivatives with constitutive expression of *mCherry*, ^55^ or *timer*^bac 39^ from the *P*_ybaJ_ promoter ^54^(chromosomal coordinates in SL1344 GenBank FQ312003.1: 528,632-528,081) and fusions of a doxycycline (DOX)-inducible promoter *P*_tetA_-*ssrB* driving expression of *P*_sifB_-*GFP-ova* coding for a degradable variant of the green fluorescent protein GFP.mut2 ^56^. All plasmids were constructed using Gibson assembly ^57^ using primers listed in Table S1 and S2.

### In-vitro *P*_*tetA*_-ssrB induction

Bacteria were grown overnight in MES-MM (100 mM MES-KOH pH 5.5, 0.4% glycerol, 15 mM NH_4_Cl, 1.5 mM K_2_SO_4_, 3 mM KH_2_PO_4_) containing 90 mg/L streptomycin and 50 mg/L kanamycin ^30^. Day cultures were inoculated at an OD_600 nm_ of 0.05 and grown for 2 h. Doxycycline (0.125 mg/L) was added to induce *P*_*tetA*_*-ssrB*. Cultures were incubated for 4 h at 37 °C with shaking (180 rpm), after which samples were withdrawn, diluted 1:10 in PBS and fluorescence intensity was measured.

### Mice

All animal experiments were approved (license 2239, Kantonales Veterinäramt Basel) and performed according to local guidelines (Tierschutz-Verordnung, Basel) and the Swiss animal protection law (Tierschutz-Gesetz). Mice (C57BL/6J *Slc11A1*^r/r 34^) were housed at 22°C (-2°C/+3°C), relative humidity of 55 +/-10%, and a 12h/12h dark/light cycle. We infected mice at 10 to 15 weeks of age and used both female and male mice which show comparable results for *Salmonella* infections ^34^. We estimated sample size by a sequential statistical design. We first infected two to three mice based on effect sizes and variation observed in our previous studies ^34,58^ and used the results to estimate group sizes for obtaining statistical significance with sufficient power.

Mice were exposed to DOX and 4% sucrose one day pre-infection in their drinking water. Mice were then infected by tail-vein injection of ∼1,200 CFU *Salmonella* grown to late-log phase in Lennox LB containing 10 mM MgCl_2_ (which increased consistency across experiments). The inoculum size was determined by plating. Intravenously infected mice show similar *Salmonella* growth rates ^39^ and *Salmonella* localization in spleen ^59^ compared to orally infected mice but exhibit less variation in *Salmonella* tissue loads between individual mice, and thus require fewer experimental animals for detecting differences with the same statistical power. For plating, mice were euthanized with carbon dioxide and spleen was homogenized in PBS containing 0.2% Triton X-100. *Salmonella* load was determined by plating on lysogeny broth agar. Activity *of P*_sifB_-*gfp-ova* and *Salmonella* survival were determined by comparing flow cytometry data and corresponding CFU counts of *Salmonella* populations. Mice were scored daily for disease signs (physical appearance: normal – 0, lack of grooming – 1, piloerection and nasal ocular discharge – 2, hunched up – 3; behavior (mobile – 0, reduced mobility – 2, immobile – 3).

### Antibodies

For depletion, purified anti-mouse CD4 (clone GK1.5) or isotype control (rat IgG2b) antibodies (obtained from Bio X Cell), a dose of 800 μg per mouse and administered intraperitoneally on days 14, 16, and 18 post-infection.

### Flow cytometry

The spleen was homogenized in 6 mL ice-cold phosphate-buffered saline containing 0.2% Triton X-100. All samples were kept on ice until and during analysis. Large host cell fragments were removed by repeated centrifugations at 100xg for 10 min at 4°C. Relevant spectral parameters were recorded in a FACS Fortessa II (Becton, Dickinson) with ‘low’ speed, using a threshold on sideward scatter (SSC) to exclude electronic noise. We used the following channels: GFP and green TIMER^bac^ component, excitation 488 nm, emission 502-525 nm (“green”); mCherry and red TIMER^bac^ component, excitation 561 nm, emission 595-664 nm (“red”) (with 633 nm laser switched off); yellow and orange autofluorescence, excitation 488 nm, emission 533-551 nm, 573-613 nm; orange autofluorescence, excitation 445 nm, emission 573-613 nm. All parameters were measured as ‘height’. Data were processed with FlowJo 10.6.1 and MATLAB R2020b. TIMER^bac^ fluorescence log color-ratios were converted into *Salmonella* division rates based on previously established calibration data ^39^.

### Quantification and statistical analysis

Statistical tests were performed with GraphPad Prism 9.3.1 as indicated in the figure legends. The respective comparisons are denoted in the Figure panels. We log-transformed CFU data to approximate normal distributions ^60^. Comparison of two groups were done by t-test. Matched measurements from the same infected mouse (such as subsets with differential promoter activity, or mutant and wild-type *Salmonella* loads) were analyzed by paired tests. Comparisons of three or more groups were done by one-way ANOVA. The impact of two factors was analyzed by two-way ANOVA. All tests were two-tailed. We deliberately avoided specifying a particular significance threshold for *P*-values (*e*.*g*., α = 0.05, 0.01, or 0.001) to avoid excessive emphasis on arbitrary cut-offs ^61^.

---

**Table 1S.**
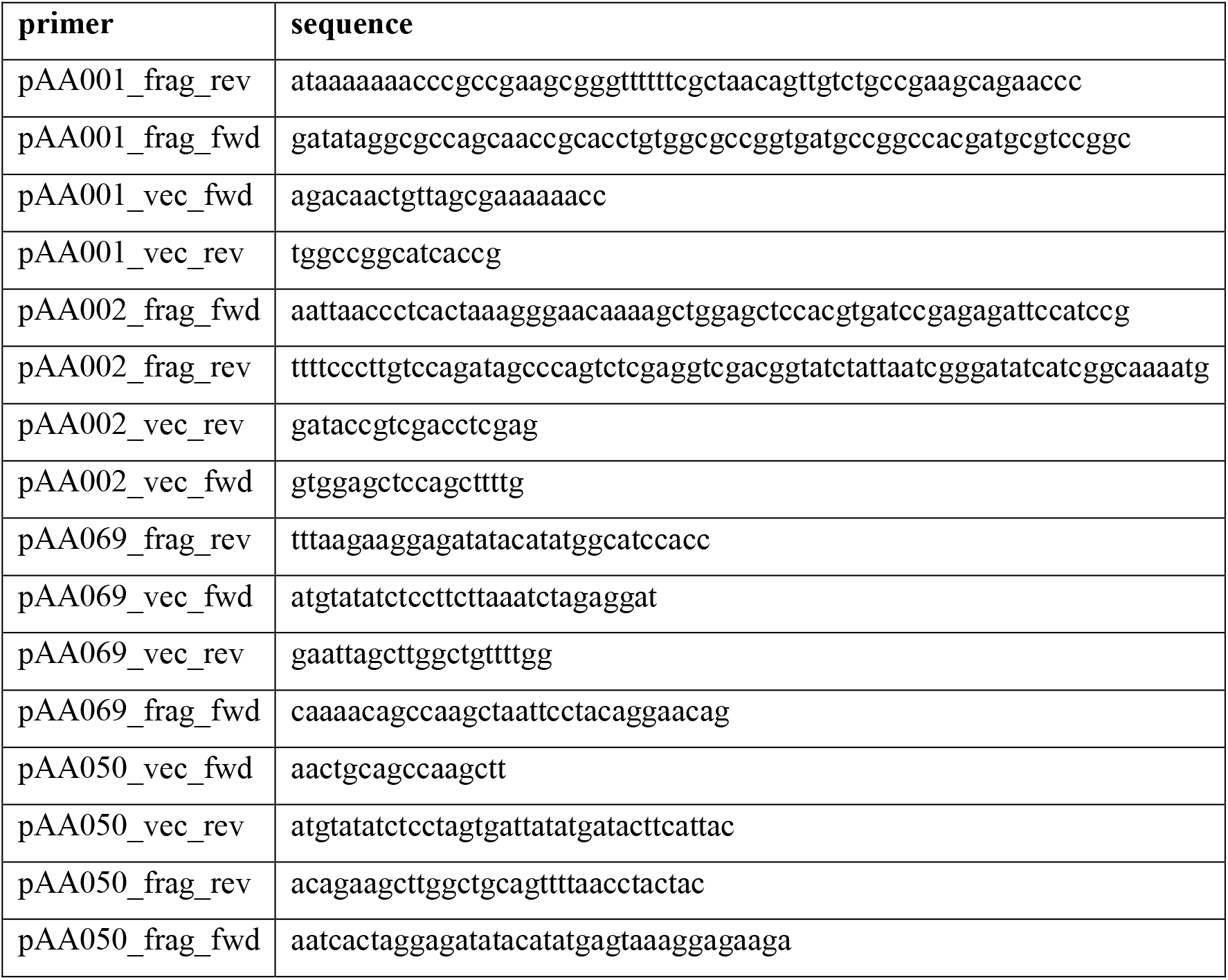

**Table 2S.**

| plasmid | Genotype |
| --- | --- |
| pAA001 | pSC101 *P<sub>tetA</sub>-ssrB P<sub>ybaJ</sub>-mCherry* (Kan<sup>R</sup>) |
| pAA002 | pFOK *ssrB*-STOP(Kan<sup>R</sup>) |
| pAA023 | pSC101 RBS (*aaagca*) *P<sub>tetA</sub>-ssrB P<sub>ybaJ</sub>-mCherry* (Kan<sup>R</sup>) |
| pAA050 | pFOK *P<sub>sifB</sub>-GFP-ova* (Kan<sup>R</sup>) |
| pAA069 | pSC101 *P<sub>tetA</sub>-ssrB P<sub>ybaJ</sub>-timer<sup>bac</sup>* (Kan<sup>R</sup>) |

## Author contributions

Outlined the study: DB and AA, performed experiments: AA, BC, SN, HA analyzed data: AA and DB, interpreted data: DB and AA, wrote manuscript: DB and AA.

## Data and materials availability

All data is available in the manuscript or the supplementary materials. Correspondence and requests for materials should be addressed to.

---

**Figure S1.**
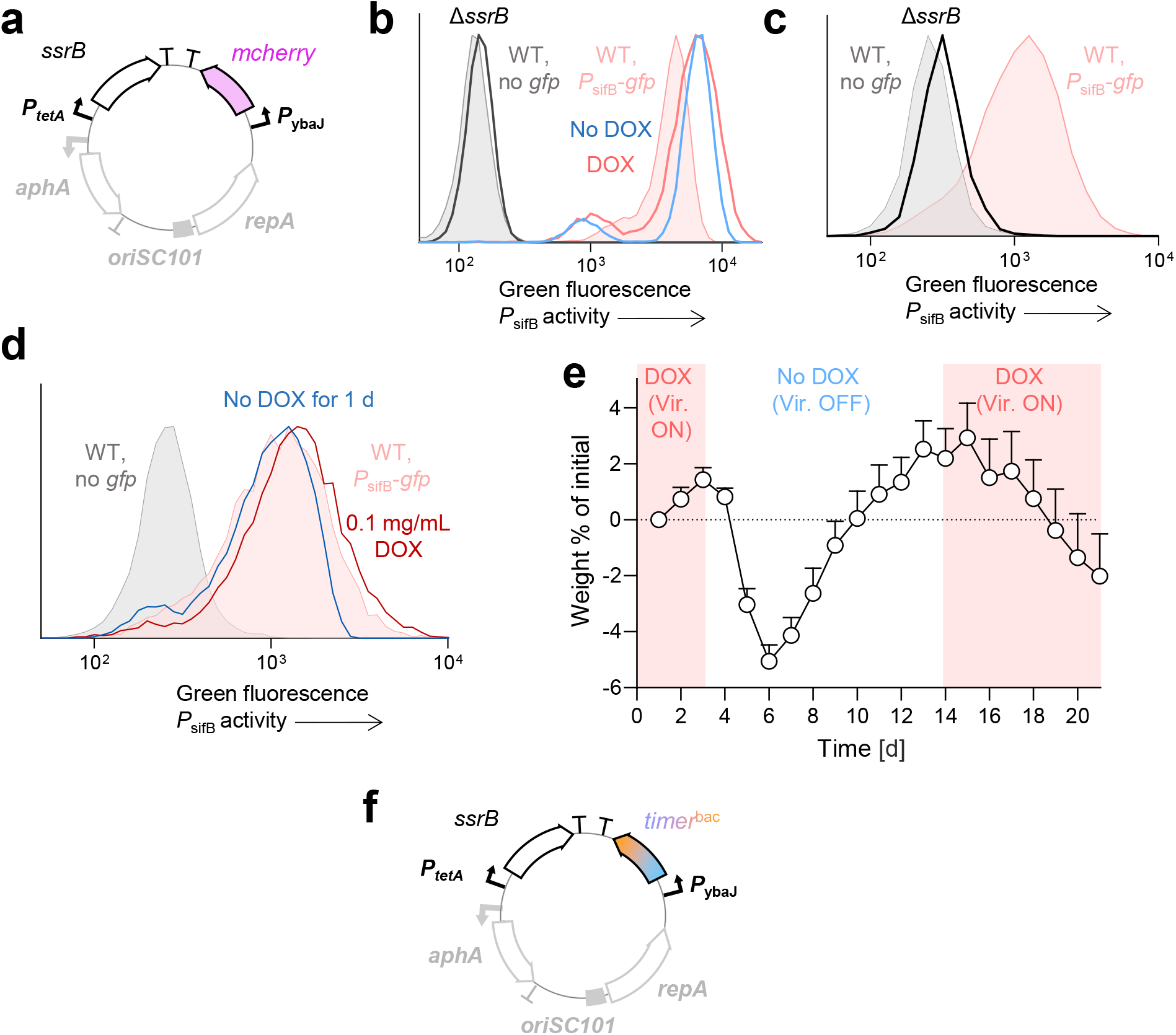
Assessment of doxycycline dosing in mice. (a) Plasmid map of the switchable-virulence construct. A constitutive *P*_*ybaJ*_*-mCherry* cassette labels all *Salmonella* irrespective of *P*_*tetA*_*-ssrB* activity. (b) DOX dependence of *P*_*sifB*_*-gfp* activity in initial not optimized construct. Autofluorescence of a (WT, wild-type) *Salmoenlla* (gray) and induced *P*_sifB_-*gfp* activity in WT is shown in (orange). Pooled data from three mice per group. (c) Green fluorescence of *P*_*sifB*_*-gfp* in *Salmonella* with inactivated *ssrB* in spleen of infected mice. Autofluorescence of a (WT, wild-type) *Salmoenlla* (gray) and induced *P*_sifB_-*gfp* activity in WT is shown in (orange). Pooled data from three mice per group. (d) Continuous *P*_*sifB*_*-gfp* activity in mice at day 6 post-infection, after receiving 0.1 mg/mL DOX for 5 days and 1 day without DOX. Autofluorescence of a (WT, wild-type) *Salmoenlla* (gray) and induced *P*_sifB_-*gfp* activity in WT is shown in (orange). Pooled data from three mice per group. (e) Body weight change (% of initial) over time. DOX was provided from day 1 to day 3, withdrawn until day 14, then re-administered until day 21. Data represent means of three to seven mice per group and their standard error of the mean. (f) Plasmid map of *P*_*tetA*_*-ssrB* with constitutive *timer*^bac^ expression.

**Figure S2.**
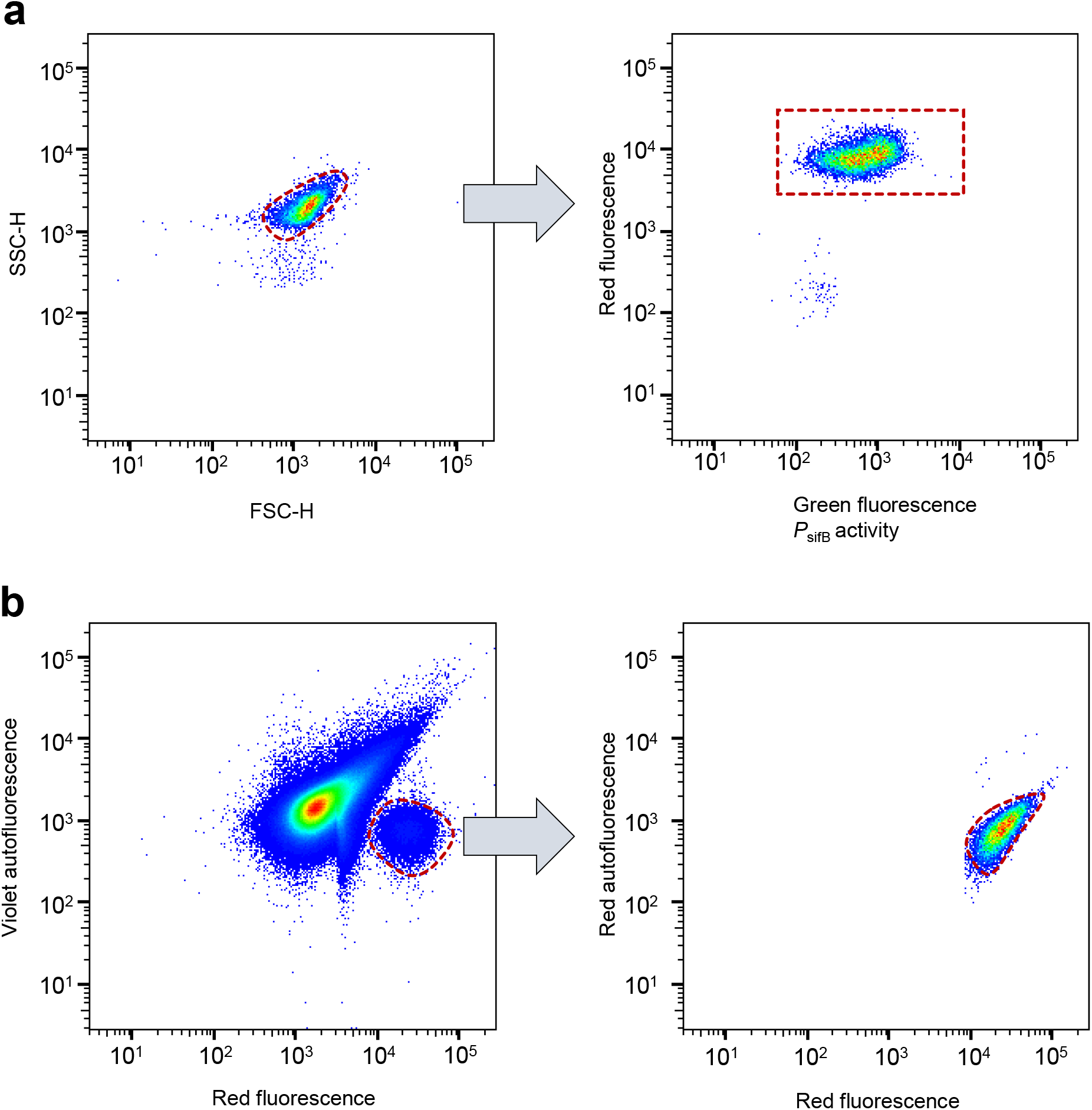
Gating strategy for in vitro and in vivo flow cytometry data. (a) Identification of single bacteria by FSC *vs* SSC, followed by gating on constitutive mCherry^+^ events to define all *Salmonella* with the construct. (b) Gate used to identify *Salmonella* with high constitutive mCherry expression in spleen homogenates.

